# Convergent evolution of increased urine concentrating ability in desert mammals

**DOI:** 10.1101/2020.09.11.294124

**Authors:** Joana L. Rocha, José C. Brito, Rasmus Nielsen, Raquel Godinho

## Abstract

One of the most celebrated textbook examples of physiological adaptations to desert environments is the unique ability that desert mammals have to produce hyperosmotic urine. Commonly perceived as an adaptation mainly observed in small rodents, the extent to which urine concentrating ability has independently evolved in distinct lineages, including medium-sized and large desert mammals, has not previously been assessed using modern phylogenetic approaches. Here, we explicitly test the general hypothesis that desert-dwelling mammals have evolved increased ability to concentrate urine compared to non-desert species, controlling for body mass and other covariates. Phylogenetic generalized least-squares models show that the mean aridity index of a species distribution range largely predicts its urine concentrating ability, even when accounting for body mass differences and phylogenetic correlations. In contrast, we find much weaker correlations between mass-adjusted basal metabolic rate and environmental variables.

## 1. Background

Deserts are defined as regions with aridity index, a ratio of annual precipitation and potential evapo-transpiration, below 0.20 [1]. Desert species independently evolved striking adaptations to cope with the water scarcity and extreme climatic and physical conditions that characterize such habitats [2–7]. For the last fifty years, this has captured the attention of many eco-physiologists who turned to desert biology to study the physiological mechanisms that help maintaining body temperature and retain water. Remarkably, most of the classic works on mammalian desert physiology, pioneered by Schmidt-Nielsen et al [8–11], have survived the test of time, as researchers have applied new perspectives and tools to test specific hypothesis regarding the evolution of adaptive traits [12,13].

Among the classical findings of physiological adaptations that minimize water loss, is the ability of some desert mammals to produce highly concentrated urine [14]. This phenotype is mostly associated with desert ‘evaders’, such as the Australian Hopping mouse, which has the record for highest hyperosmotic urine (above 9000 mOsm/kg) [3,15]. Evaders, part of a classification system proposed by Willmer *et al*. (2000), are small body-sized animals that are able to evade extreme conditions through behavior [5]. By contrast, large-sized mammals unable to shelter from extreme climates that are forced to withstand heat, are called ‘endurers’, and medium-sized mammals unable to evade nor withstand extremes as efficiently as evaders and endurers are called ‘evaporators’ [5]. Because mammalian urine concentrating ability (mOsm/Kg) is negatively correlated with body mass [16], desert evaders stand out in this capacity [15]. However, when correcting for differences in body mass, the capacity to highly concentrate urine seems to have independently evolved in deserts. For example, evaporators such as the fennec fox can concentrate urine up to 4022 mOsm/Kg, which is much more concentrated than in similar-sized non-desert counterparts and, when mass-adjusted, nearly as impressive as the 5500 mOsm/Kg of a lesser Egyptian gerbil [5,17,18].

Classical comparative physiology studies have provided evidence for convergent adaptive evolution of ecologically relevant phenotypes in multiple desert mammals. A textbook example includes basal metabolic rate (BMR), which is positively correlated with zoogeographical zones, even when accounting for phylogeny and body mass [19,20]. Desert mammals evolved lower production of metabolic heat (low BMR) to maintain body temperatures and avoid water loss by evaporative cooling [19–23]. Surprisingly, and to the best of our knowledge, no study to date has attempted to identify continuous climatic or environmental variables that help explain variation in urine concentrating ability, one of the most celebrated examples of mammalian desert adaptation. Moreover, and in contrast to BMR, previous studies concerning urine concentrating ability have ignored phylogenetic correlations when analyzing associations between this trait and the environment [16,24].

Here, we explicitly test the hypothesis that the aridity index of a species range predicts maximum urine concentrating ability, even when accounting for body mass and phylogenetic relationship, and provide statistical evidence that the ability to avoid water loss by producing hyperosmotic urine has evolved independently in multiple phylogenetic lineages of desert mammals. Additionally, we use similar models to re-analyze mass adjusted BMR for species in which urine osmolality was obtained, and test aridity index and temperature variables as predictors for metabolism in deserts.

## 2. Methods

Phenotypes were obtained by extensive literature search and by taking advantage of large datasets from previously published revision studies [16,19,24,25]. A total of 108 mammalian species’ maximum urine concentrating abilities came from two previous studies by Beuchat [16,24], which recorded the specific condition (factor defined as C) under which urine osmolality (hereafter mOsm) was measured, including: dehydrated (D), given salt (S) or protein (P) loading, combinations of these conditions (DP, DS, SP, DSP), or treatment not specified (X), as well as the method used to measure urine mOsm (see 15,23 for more information). We also recorded the highest urine osmolality values for 13 additional species from other studies [18,26–30], and also noted the experimental conditions (C) the study-cases were subjected to, the method (M) used to measure urine mOsm, and the study-species diet (Di), following the same notation as Beuchat (see 15,23 for more information). For species with more than one record of urine mOsm, due to different treatments/studies, we opted to keep values taken from captive study-subjects under controlled/experimentally induced dehydration, as opposed to measurements of unknown condition taken from the field. Ultimately, only one individual was used per species.

As mOsm is a plastic trait, the experimental conditions under which this parameter is measured can greatly affect estimates. In particular, the degree of hydration of the study animals naturally affects urine osmolality. As such, we curated two datasets: 1) a larger dataset including all 121 reported measurements of mOsm from Beuchat [16,24] and our search [18,26–30]; 2) a subset of the larger dataset only containing 87 observations in which the study-subject, in captivity, was dehydrated (D) and not given food or any other treatment. However, we note that there is still some residual variation in the number of days under water deprivation for an animal to be considered fully dehydrated and variation in the method used for measuring mOsm. Also, for the 121 species dataset, 24 records came from studies that did not report the procedures used to measure mOsm. A full detail of the conditions and studies can be found in electronic Supplementary Material TableS1. Downstream analyses were performed on both data sets.

The distribution polygons of each species for which physiological data were available were downloaded from the IUCN database [31,32]. We also downloaded the following environmental variables at about 1km spatial resolution: annual aridity index (AI) from CGIAR-CSI Global Aridity database [33], and annual mean temperature (BIO1), maximum temperature of the warmest month (BIO5), maximum temperature of the coldest month (BIO6), and temperature annual range (BIO7) from the WorldClim database [34]. We then used a Geographical Information System to calculate mean values for each environmental variable within the distribution polygon of each species. For species with global distributions ranges, encompassing humid to arid environments, and for which phenotypes under review were sampled from a specific region, we partitioned the original distribution polygon accordingly and calculated mean values for the sampled zone. To avoid the effects of collinearity we tested for correlations between pairs of variables using a Spearman correlation test before testing them together as predictors for mOsm and BMR in downstream analyses. We used a previously published mammalian tree estimated using both nuclear and mitochondrial genes [35] and pruned the tree to only retain the species included in or study. For visualization of trait evolution on the tree, we projected states for maximum mOsm and bioclimatic variables, estimated using maximum likelihood, onto the internal edges and nodes of the tree using a color gradient. The estimation was done using the *fastAnc* option in the ‘contMap’ function implemented in ‘phytools’ [36].

Phylogenetic generalized least squares (PGLS) as implemented in ‘phytools’ were performed using the ‘gls’ function with the model of Pagel (1999) [37] to test different linear models with increasing complexity. Briefly, the model of Pagel adjusts the off-diagonal elements of variance-covariance matrix in a Brownian evolution model with a multiplicative factor, λ, such that when λ = 0 a star phylogeny is obtained representing no phylogenetic signal, while when λ = 1 a standard Brownian on the reference tree is obtained. As residuals of maximum urine osmolality are not normally distributed (**Figure S1**), we performed PGLS analyses on log10(maximum mOsm) using log10(body mass) as covariate. Mean annual aridity index (hereafter mean AI) and uncorrelated mean temperature variables were used in PGLS models as potential predictors of urine concentrating ability. Condition (C), method (M) and diet (Di) were also included as covariates. For the dataset including only dehydrated individuals, the same PGLS models were tested with AI converted to a categorical variable using the following binning defined in [33]: humid for AI > 0.65, dry sub-humid for 0.5< AI < 0.65, semi-arid for 0.2 < AI < 0.5, and arid for AI < 0.2. To test for phylogenetic signal for specific traits, we calculated Pagel’s λ on urine concentrating ability, body mass and AI using the function ‘phylosig’ as implemented in ‘phytools’.

## 3. Results

We analyzed a dataset with information on maximum urine concentrating abilities, mean annual aridity index and WorldClim temperatures for a total of 121 species of mammals, with available DNA sequence information, spanning 6 g to 479 kg of body mass. Using the classification of [33] the dataset contained 32, 35, 15, and 39 species with distribution ranges classified as arid, semi-arid, dry sub-humid, and humid, respectively. Ancestral state reconstruction across our mammalian dataset’s phylogeny shows that the ability to produce hyperosmotic urine has evolved multiple times in species with distribution ranges characterized by low mean AI (**Figure 1**).

**Figure 1.**
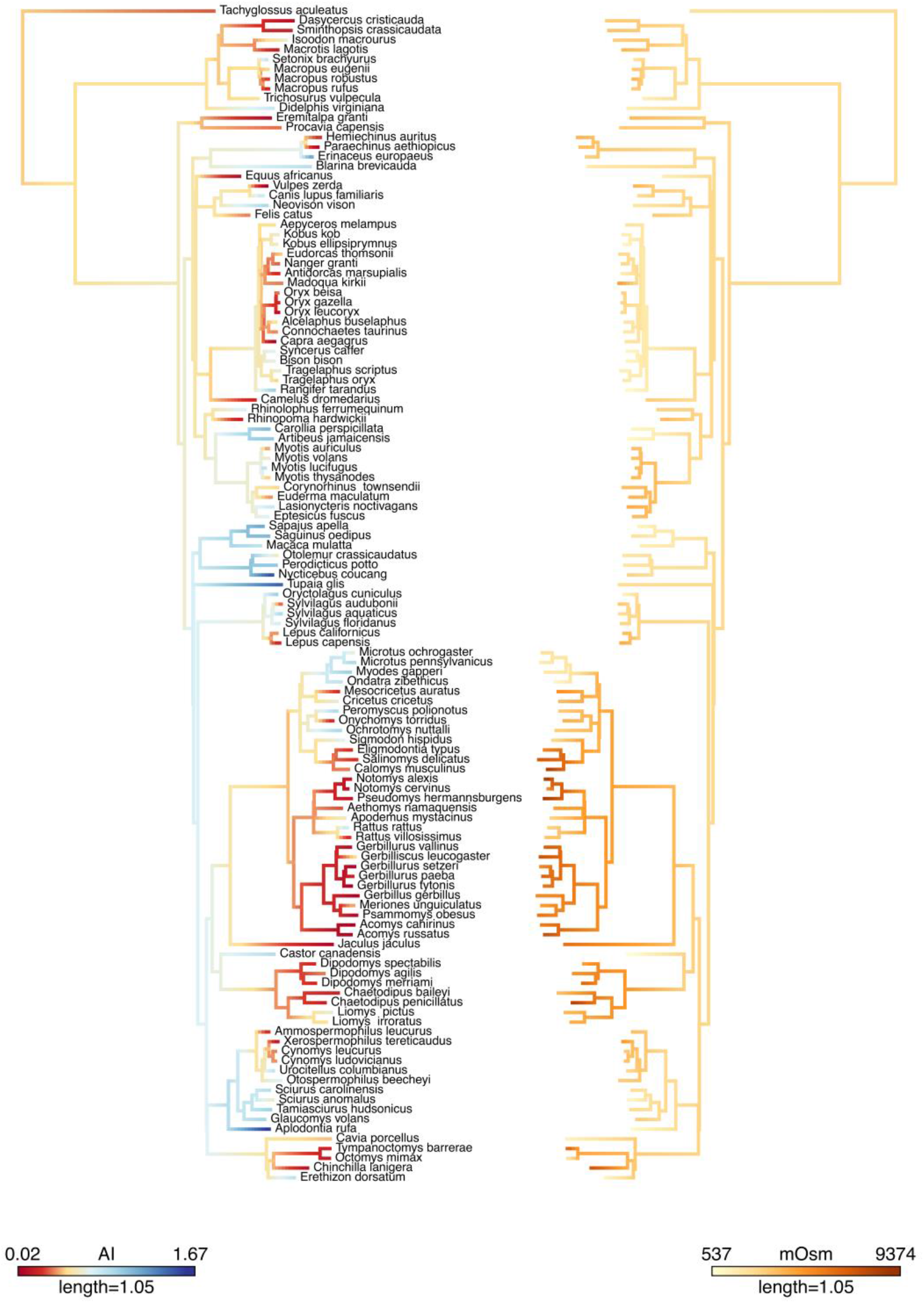
Ancestral state reconstruction for mean annual aridity index (AI, left) and maximum urine osmolalities (mOsm/kg; right) in a phylogenetic tree with 121 mammalian species.

When testing for phylogenetic signal using Pagel’s λ we found highly significant phylogenetic signal for most variables, validating the relevance of adjusting the linear models for phylogenetic covariance (**Table 1**). The mean AI correlates with all temperature variables, except mean annual temperature range (**Figure S3**). To avoid collinearity, all WorldClim temperatures but the latter were excluded from PGLS analysis with mean AI (**Table S2**).

**Table 1.**
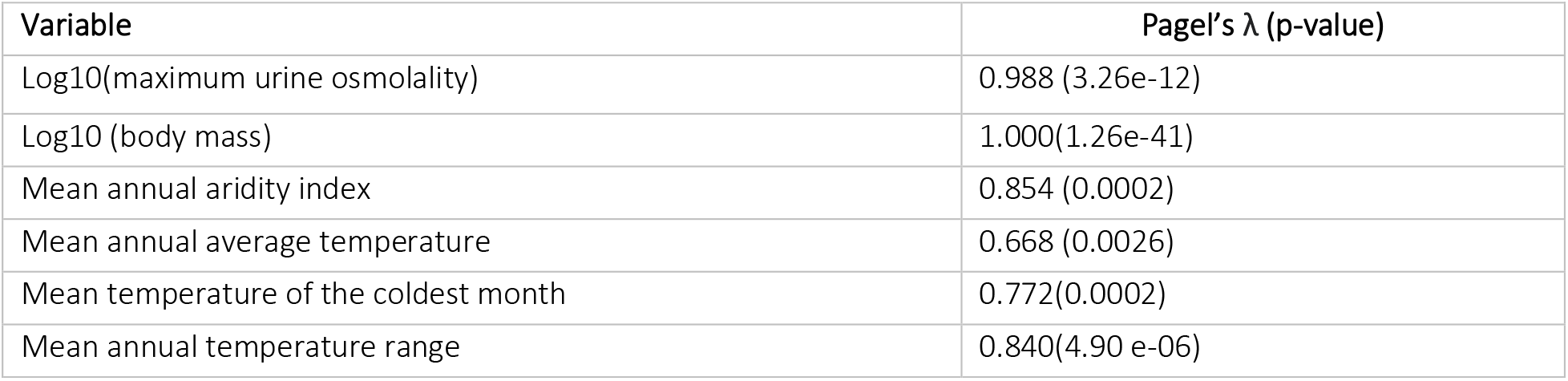
Phylogenetic signal (λ) for variables with significant p-value. Variables for which λ is close to 1, and with p-values < 0.05, are variables with significant phylogenetic signals.

All PGLS models tested in this study point to mean annual aridity index as a highly significant and important predictor of mammalian maximum urine concentration ability (**Table 2, TableS2**). The lower the mean AI, the more efficient mammals are able to concentrate urine (**Figure 2, FigureS2**), even when accounting for the differences in body mass, diet, experimental condition, methodology and phylogeny (**Table 2**). Overall, species inhabiting arid and semi-arid environments can maximize urine concentration well above the levels of dry sub-humid and humid counterparts (**Figure 2**). In congruence with past studies, our results also show that log-transformed body mass is also negatively correlated with log-transformed maximum urine osmolality [16], though not to the same extent as mean aridity index (**Table 2**, **Figure S2**). Diet is also a significant predictor of maximum urine osmolality in fructivorous mammals, which significantly concentrate more urine than strictly carnivore species (**Table 2**). Overall, no strong correlation was found between urine concentrating ability and temperature variables (**Figure S4, TableS2**).

**Table 2.** PGLS models for predicting mammalian log10(maximum urine osmolality). Model notation refers to the combination of variables used as predictors while taking phylogeny into account. Highlighted in bold are significant p-values. Variables are AI = mean aridity index, BM = Log10 body mass (kg), C = condition (dehydrated-D relative to fed-F, given salt load-S, given protein load-P, unknown-x and combinations of these), M = Method (unknown-? relative to addition of solutes-A, Freezing point-F, Vapor pressure-V, and combinations of these), Di = Diet (Carnivorous-C relative to Fructivorous-Fr, Granivorous-G, Herbivorous-H, Insectivorous-I, Omnivorous-O, and combinations of these).

| *Dataset* | *Model* | *AIC* | *variable* | *Coefficient* | *s.e* | *t-value* | *p-value* |
| --- | --- | --- | --- | --- | --- | --- | --- |
| *All*<br>(*n*=121) | AI + BM + C + M +Di | -16.3 | intercept | 3.703 | 0.097 | 37.99 | **0.0000** |
|  |  |  | AI | -0.368 | 0.043 | -8.47 | **0.0000** |
|  |  |  | BM | -0.082 | 0.015 | -5.54 | **0.0000** |
|  |  |  | DF<sub>condition</sub> | 0.023 | 0.080 | 0.28 | 0.7763 |
|  |  |  | DP<sub>condition</sub> | -0.081 | 0.108 | -0.75 | 0.4537 |
|  |  |  | DPS<sub>condition</sub> | 0.020 | 0.178 | 0.11 | 0.9104 |
|  |  |  | DS<sub>condition</sub> | -0.047 | 0.106 | -0.45 | 0.6573 |
|  |  |  | P<sub>condition</sub> | -0.006 | 0.077 | -0.08 | 0.9341 |
|  |  |  | S<sub>condition</sub> | 0.049 | 0.065 | 0.75 | 0.4536 |
|  |  |  | SP<sub>condition</sub> | 0.148 | 0.148 | 1.00 | 0.3202 |
|  |  |  | X<sub>condition</sub> | -0.049 | 0.047 | -1.06 | 0.2911 |
|  |  |  | A<sub>method</sub> | -0.068 | 0.153 | -0.44 | 0.6592 |
|  |  |  | F<sub>method</sub> | 0.016 | 0.039 | 0.41 | 0.6801 |
|  |  |  | FV<sub>method</sub> | 0.058 | 0.097 | 0.60 | 0.5530 |
|  |  |  | V<sub>method</sub> | 0.106 | 0.055 | 1.93 | 0.0570 |
|  |  |  | Cl<sub>diet</sub> | -0.158 | 0.141 | -1.12 | 0.2649 |
|  |  |  | Fr<sub>diet</sub> | -0.533 | 0.149 | -3.57 | **0.0005** |
|  |  |  | G<sub>diet</sub> | 0.013 | 0.103 | 0.13 | 0.8980 |
|  |  |  | H<sub>diet</sub> | -0.132 | 0.088 | -1.50 | 0.1366 |
|  |  |  | I<sub>diet</sub> | -0.130 | 0.107 | -1.22 | 0.2260 |
|  |  |  | O<sub>diet</sub> | -0.055 | 0.095 | -0.58 | 0.5634 |
| *Dehydrated*<br>(*n*=87) | AI + BM + M +Di | -34.7 | intercept | 3.679 | 0.132 | 27.83 | **0.0000** |
|  |  |  | AI | -0.344 | 0.043 | -7.96 | **0.0000** |
|  |  |  | BM | -0.067 | 0.016 | -4.18 | **0.0001** |
|  |  |  | A<sub>method</sub> | -0.107 | 0.141 | -0.75 | 0.4528 |
|  |  |  | F<sub>method</sub> | -0.007 | 0.042 | -0.16 | 0.8702 |
|  |  |  | FV<sub>method</sub> | 0.096 | 0.104 | 0.93 | 0.3563 |
|  |  |  | V<sub>method</sub> | 0.029 | 0.070 | 0.42 | 0.6758 |
|  |  |  | Cl<sub>diet</sub> | -0.098 | 0.165 | -0.60 | 0.5536 |
|  |  |  | Fr<sub>diet</sub> | -0.406 | 0.178 | -2.28 | **0.0254** |
|  |  |  | G<sub>diet</sub> | 0.090 | 0.142 | 0.63 | 0.5281 |
|  |  |  | H<sub>diet</sub> | -0.106 | 0.135 | -0.79 | 0.4333 |
|  |  |  | I<sub>diet</sub> | -0.049 | 0.140 | -0.35 | 0.7261 |
|  |  |  | O<sub>diet</sub> | 0.002 | 0.139 | 0.02 | 0.9873 |

**Figure 2.**
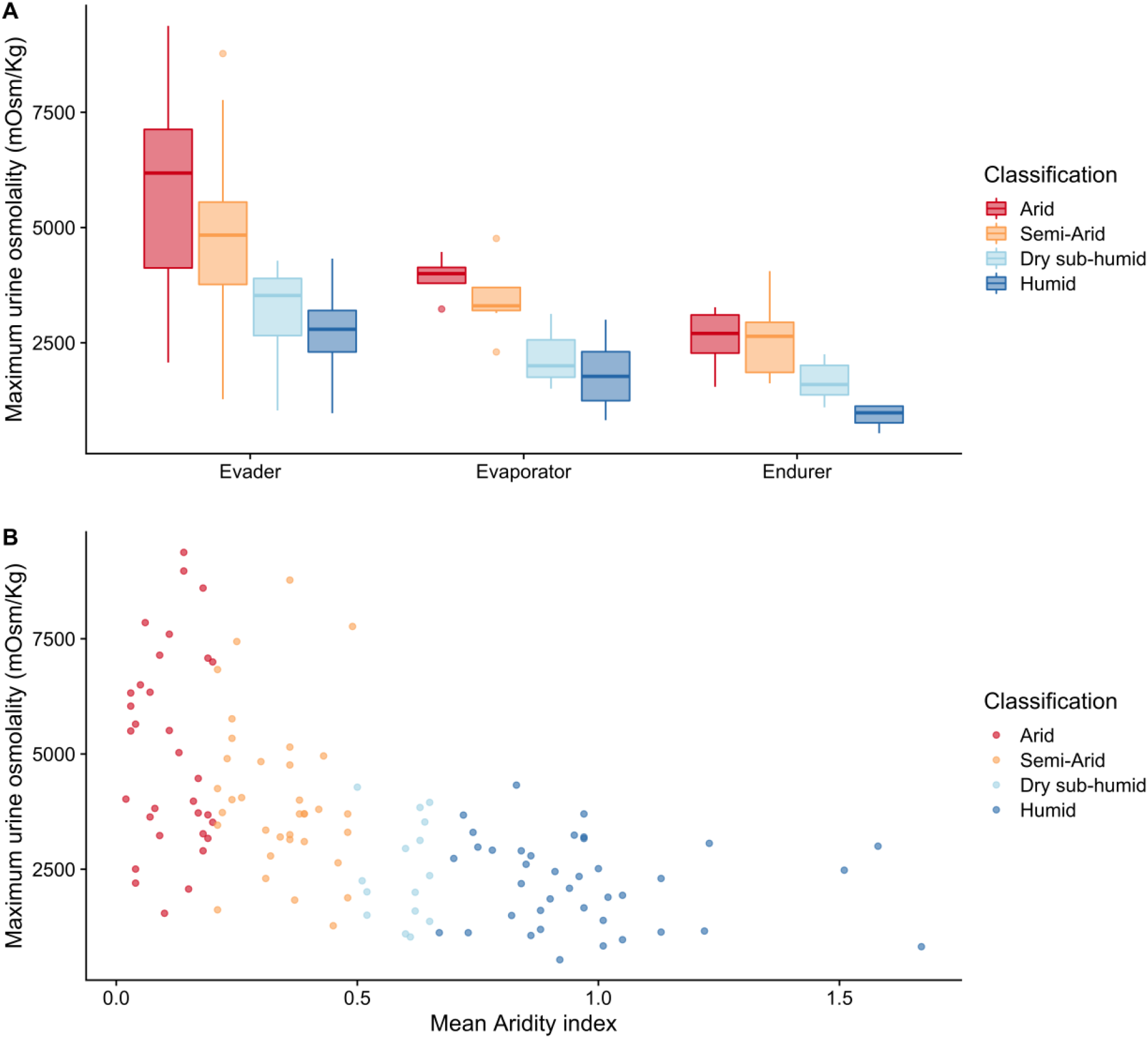
Maximum urine osmolality (mOsm/Kg) and mean aridity index (AI). (A) Boxplot for maximum urine osmolality in endurers, evaders, and evaporators from environments with different degree of aridity, based on 121 different species. Endurers, evaders, and evaporators are classified based on body mass following [5] and aridity classification according to [33]. (B) Scatterplot using mean aridity index as a continuous variable.

When the experimental condition was used as co-variate in the PGLS analysis, its role in predicting maximum urine osmolality was not significant (**Table 2, Table S2**). Nonetheless, we also tested similar PGLS models on a subset of the data that only included individuals fulfilling the dehydrated criterion (see Methods section), resulting in a smaller dataset including 87 species (18 arid, 26 semi-arid, 13 dry sub-humid, and 30 humid). The results from this analysis are similar to those of the larger dataset conditions (**Table 2**). When using AI as a discrete variable we found that urine concentrating ability is significantly higher in species from arid environments than in species from semi-arid, dry sub-humid and humid regions (**Table S3**). In all analyses, the aridity index of the species distribution is a stronger predictor of maximum urine osmolality than body mass and diet. Notably, experimental condition and methodology are non-significant.

We also obtained mass-adjusted basal metabolic rates (BMR) for 84 of the 121 species from our dataset. BMR is weakly correlated with bioclimatic variables (**Figure S5**), and differences in this parameter are better explained by log10 body mass (**TableS4**).

## 4. Discussion

The extent to which different mammalian species from desert environments have significantly higher urine concentrating abilities, a phenotype to better retain water in extreme conditions of water-deprivation and extreme temperature ranges, has not previously been subject to rigorous statistical analyses that account for phylogeny. Here we show that increased ability to concentrate urine, measured as maximum urine osmolality, has convergently evolved in many desert mammals (**Figure 1**).

PGLS analysis on a multi-species dataset revealed that desert-dwelling species effectively concentrate more urine than non-desert species, even when accounting for both body mass and ancestry, and furthermore, that differences in mammalian maximum urine concentrating ability can be predicted, to some degree, by the aridity of species’ range. Our measurement of environmental aridity is inherently imprecise as it does not account for spatial and temporal changes in the environmental conditions nor in the species distributions, and it implicitly assumes a uniform density distribution across the species range [33,34]. Still, we find it more rigorous than the often-used binary approach of classifying species as ‘mesic’ or ‘xeric’ based on eye-assessment of species distribution range or habitat descriptions (e.g. [19,24]). Despite these challenges, and while acknowledging additional noise added by varying experimental conditions and methodologies, we find a strongly significant correlation between maximum urine osmolality and mean annual aridity index. This observation is similar to what has been previously reported in other desert adaptive phenotypes such as basal metabolic rate [19,20]. However, we find a much weaker correlation for BMR and environmental variables when analyzing 84 of the same species as in the mOsm analysis, suggesting that, in contrast to our results for mOsm, differences in BMR among species are mostly explained by body mass differences and are only weakly predicted by environmental variables [38].

A contentious debate in the fields of ecological and evolutionary physiology is whether phenotypic differences between species and populations inhabiting contrasting environments are due to genetic (evolutionary) adaptations or due to plastic responses, and difference in the ability to concentrate urine is no exception. Although all mammals, when water-deprived, are able to increase urine concentration in relation to their optimal hydrated states [14,39], our results for a dataset in which species were all captive and water-deprived, provide strong evidence that increased maximal urine concentration is an adaptation to high aridity and has evolved multiple times in different phylogenetic lineages. The hypothesis that higher maximum urine concentrating ability in mammals is an adaptive trait is also supported by an observed increase in renal medullary thickness in desert mammals [3,16].

Our PGLS analyses also suggest that, among the many physical and climatic challenges faced by desert species, aridity, more than temperature, has been one of the main selective pressures increasing maximum urine concentrating ability, and driving its repeated evolution in different desert mammalian lineages. As population and comparative genomics studies in desert environments continue to search for the genetic basis of desert adaptation (e.g. [40–43]), maximum urine concentrating ability might be one of the primary physiological traits that lends itself to more detailed genetic analyses.

## Author’s contributions

J.L.R contributed to the conception and design of the study, data collection, analysis, and drafted the manuscript. R.G, J.C.B, R.N contributed to the study’s conception and design, interpretation of data, editing and critical revision. All authors approve publication and agree to be accountable for all aspects of the work.

## Acknowledgments

We thank Pedro Tarroso and Antigoni Kaliontzopoulou for discussions that helped conceiving this study.

## Ethics

No ethical approval was required as this work only uses publicly available data.

