## Supplemental Table S1 for "Convergent evolution of increased urine concentrating ability in desert mammals": Rocha_et_al_SupplementaryMaterial_TableS1.html

**Table S1- Data
collection from past studies concerning 121 species listed alphabetically by
scientific name, followed by family and order.**
For each species, the following variables are reported: category according to
the endurer-evader-evaporator concept, C = condition  (dehydrated-D, fed-F,
given salt load-S, given protein load-P, unknown-x and combinations of these),
M = Method (unknown-?, addition of solutes-A,
Freezing point-F, Vapor pressure-V, and combinations of these), Di = Diet
(Carnivorous-C, Fructivorous-Fr, Granivorous-G, Herbivorous-H, Insectivorous-I,
Omnivorous-O, and combinations of these), BM = body mass (kg), mOsm =
maximum urine osmolality (mosmol/Kg H_2_O), cAI = mean aridity index
(categorical: Arid, Semi-Arid, Dry sub-humid, Humid), AI = mean annual aridity
index (continuous) , BIO1 = mean Annual Mean Temperature, BIO5= mean Maximum
Temperature of Warmest Month, BIO6 = mean Minimum Temperature of Coldest Month,
BIO7 = mean Temperature Annual Range (BIO5-BIO6). Mass-adjusted Basal Metabolic
Rate (ml of O_2_ per g per day) are also reported.  References for mOsm and BMR are also reported
after the measurement.

|  |  |  |  |  |  |  |  |  |  |  |  |  |  |
| --- | --- | --- | --- | --- | --- | --- | --- | --- | --- | --- | --- | --- | --- |
| **Species\_Family\_Order** | **Category** | **C** | **M** | **Di** | **BM** | **mOsm** | **cAI** | **AI** | **BIO1** | **BIO5** | **BIO6** | **BIO7** | **BMR** |
| Acomys\_cahirinus\_MURIDAE\_RODENTIA | Evader | D | V | O | 0.033 | 6039[1] | Arid | 0.03 | 23.3 | 37.99 | 7.45 | 30.54 | 1.1[2,3] |
| Acomys\_russatus\_MURIDAE\_RODENTIA | Evader | x | ? | O | 0.053 | 7850[4] | Arid | 0.06 | 23.99 | 36.9 | 9.98 | 26.92 | 0.77[2,3] |
| Aepyceros\_melampus\_BOVIDAE\_CETARTIODACTYLA | Endurer | D | F | H | 38 | 2250[1] | Dry sub-humid | 0.51 | 21.99 | 31.46 | 10.8 | 20.66 |  |
| Aethomys\_namaquensis\_MURIDAE\_RODENTIA | Evader | D | V | O | 0.107 | 4835[1] | Semi-Arid | 0.3 | 19.81 | 31.6 | 5.41 | 26.19 | 0.84[2,3] |
| Alcelaphus\_buselaphus\_BOVIDAE\_CETARTIODACTYLA | Endurer | D | F | H | 88 | 2010[1] | Dry sub-humid | 0.52 | 23.27 | 33.59 | 11.94 | 21.65 |  |
| Ammospermophilus\_leucurus\_SCIURIDAE\_RODENTIA | Evader | D | F | O | 0.093 | 3730[1] | Semi-Arid | 0.22 | 11.73 | 32.64 | -5.39 | 38.03 | 0.98[2,3] |
| Antidorcas\_marsupialis\_BOVIDAE\_CETARTIODACTYLA | Endurer | x | F | H | 34 | 1620[4] | Semi-Arid | 0.21 | 19.41 | 31.92 | 4.16 | 27.77 |  |
| Aplodontia\_rufa\_APLODONTIIDAE\_RODENTIA | Evaporator | D | F | H | 0.785 | 820[1] | Humid | 1.67 | 7.96 | 24.34 | -3.26 | 27.59 | 0.52[2,3] |
| Apodemus\_mystacinus\_MURIDAE\_RODENTIA | Evader | x | ? | O | 0.036 | 2950[4] | Dry sub-humid | 0.6 | 11.17 | 28.89 | -3.7 | 32.59 | 1.67[2,3] |
| Artibeus\_jamaicensis\_PHYLLOSTOMIDAE\_CHIROPTERA | Evader | D | V | Fr | 0.06 | 972[1] | Humid | 1.05 | 23.97 | 31.99 | 15.76 | 16.23 | 1.25[2,3] |
| Bison\_bison\_BOVIDAE\_CETARTIODACTYLA | Endurer | D | ? | H | 353 | 1124[1] | Humid | 0.67 | -2.04 | 22.08 | -26.12 | 48.2 |  |
| Blarina\_brevicauda\_SORICIDAE\_EULIPOTYPHLA | Evader | D | F | O | 0.019 | 1857[1] | Humid | 0.9 | 6.61 | 27.23 | -14.89 | 42.11 | 3.2[2,3] |
| Calomys\_musculinus\_CRICETIDAE\_RODENTIA | Evader | x | V | O | 0.0127 | 8773[5] | Semi-Arid | 0.36 | 15.27 | 29.24 | 1.96 | 27.28 | 1.63[2,3] |
| Camelus\_dromedarius\_CAMELIDAE\_CETARTIODACTYLA | Endurer | D | F | H | 479 | 3170[1] | Arid | 0.19 | 22.51 | 37.81 | 6.91 | 30.9 | 0.1[2,3] |
| Canis\_lupus\_familiaris\_CANIDAE\_CARNIVORA | Evaporator | x | F | C | 15 | 2608[1] | Humid | 0.85 | 0.18 | 22.62 | -21.02 | 43.64 | 0.33[2,3] |
| Capra\_aegagrus\_BOVIDAE\_CETARTIODACTYLA | Endurer | D | ? | H | 23 | 2200[1] | Arid | 0.04 | 13.93 | 32.96 | -3.39 | 36.34 |  |
| Carollia\_perspicillata\_PHYLLOSTOMIDAE\_CHIROPTERA | Evader | D | V | Fr | 0.015 | 1137[1] | Humid | 1.13 | 24.36 | 31.52 | 16.84 | 14.68 | 2.11[2,3] |
| Castor\_canadensis\_CASTORIDAE\_RODENTIA | Endurer | x | F | H | 25 | 537[1] | Humid | 0.92 | 3.22 | 24.2 | -17.29 | 41.49 |  |
| Cavia\_porcellus\_CAVIIDAE\_RODENTIA | Evader | D | F | H | 0.75 | 1275[1] | Semi-Arid | 0.45 | 12.1 | 21.34 | 0.93 | 20.42 | 0.55[2,3] |
| Chaetodipus\_baileyi\_HETEROMYIDAE\_RODENTIA | Evader | DS | ? | G | 0.025 | 3680[1] | Arid | 0.19 | 20.28 | 37.93 | 3.26 | 34.67 | 1.31[2,3] |
| Chaetodipus\_penicillatus\_HETEROMYIDAE\_RODENTIA | Evader | x | F | G | 0.017 | 8600[1] | Arid | 0.18 | 19.65 | 37.93 | 2.71 | 35.22 | 1.31[2,3] |
| Chinchilla\_lanigera\_CHINCHILLIDAE\_RODENTIA | Evader | D | F | H | 0.475 | 7599[1] | Arid | 0.11 | 12.93 | 22.62 | 4.26 | 18.36 | 1.4[2,3] |
| Connochaetes\_taurinus\_BOVIDAE\_CETARTIODACTYLA | Endurer | D | ? | H | 200 | 1832[1] | Semi-Arid | 0.37 | 21.08 | 32.2 | 6.64 | 25.56 | 0.7[2,3] |
| Corynorhinus\_townsendii\_VESPERTILIONIDAE\_CHIROPTERA | Evader | D | F | I | 0.008 | 3700[1] | Semi-Arid | 0.48 | 11.32 | 30.39 | -5.24 | 35.63 | 0.29[2,3] |
| Cricetus\_cricetus\_CRICETIDAE\_RODENTIA | Evader | D | F | O | 0.108 | 3840[1] | Dry sub-humid | 0.63 | 4.93 | 26.17 | -14.75 | 40.92 | 0.64[2,3] |
| Cynomys\_leucurus\_SCIURIDAE\_RODENTIA | Evaporator | D | F | H | 0.962 | 3145[1] | Semi-Arid | 0.36 | 5.54 | 27.84 | -13.1 | 40.94 |  |
| Cynomys\_ludovicianus\_SCIURIDAE\_RODENTIA | Evaporator | D | F | H | 1.145 | 3704[1] | Semi-Arid | 0.39 | 10.84 | 31.64 | -9.08 | 40.72 | 0.38[2,3] |
| Dasycercus\_cristicauda\_DASYURIDAE\_DASYUROMORPHIA | Evaporator | D | A | O | 3 | 3231[1] | Arid | 0.09 | 22.36 | 38.3 | 5.64 | 32.66 | 0.52[2,3] |
| Didelphis\_virginiana\_DIDELPHIDAE\_DIDELPHIMORPHIA | Evaporator | DP | F | O | 5 | 1497[1] | Humid | 0.82 | 15.9 | 31.59 | 0.07 | 31.52 | 0.33[2,3] |
| Dipodomys\_agilis\_HETEROMYIDAE\_RODENTIA | Evader | D | ? | G | 0.061 | 5150[1] | Semi-Arid | 0.36 | 14.34 | 31.37 | 0.88 | 30.5 | 1.05[2,3] |
| Dipodomys\_merriami\_HETEROMYIDAE\_RODENTIA | Evader | DP | V | G | 0.035 | 6832[1] | Semi-Arid | 0.21 | 18.08 | 35.21 | 1.42 | 33.79 | 1.18[2,3] |
| Dipodomys\_spectabilis\_HETEROMYIDAE\_RODENTIA | Evader | D | F | G | 0.1 | 4900[4] | Semi-Arid | 0.23 | 15.21 | 33.46 | -2.8 | 36.26 |  |
| Eligmodontia\_typus\_CRICETIDAE\_RODENTIA | Evader | D | V | G | 0.02 | 5763[1) | Semi-Arid | 0.24 | 7.75 | 18.54 | -5.8 | 24.34 |  |
| Eptesicus\_fuscus\_VESPERTILIONIDAE\_CHIROPTERA | Evader | D | F | I | 0.013 | 3675[1] | Humid | 0.72 | 10.29 | 28.84 | -7.92 | 36.76 | 2.42[2,3] |
| Equus\_africanus\_EQUIDAE\_PERISSODACTYLA | Endurer | D | F | H | 129 | 1545[4] | Arid | 0.1 | 27.48 | 37.16 | 18.32 | 18.84 |  |
| Eremitalpa\_granti\_CHRYSOCHLORIDAE\_AFROSORICIDA | Evader | D | F | I | 0.027 | 3820[4] | Arid | 0.08 | 18.81 | 29.88 | 8.13 | 21.75 | 0.49[2,3] |
| Erethizon\_dorsatum\_ERETHIZONTIDAE\_RODENTIA | Evaporator | P | F | H | 15 | 1195[1] | Humid | 0.88 | 1.74 | 23.46 | -19.38 | 42.84 | 0.25[2,3] |
| Erinaceus\_europaeus\_ERINACEIDAE\_EULIPOTYPHLA | Evader | P | F | O | 0.637 | 3062[1] | Humid | 1.23 | 7.34 | 22.56 | -5.18 | 27.74 | 0.4[2,3] |
| Euderma\_maculatum\_VESPERTILIONIDAE\_CHIROPTERA | Evader | D | F | I | 0.016 | 4000[1] | Semi-Arid | 0.38 | 9.79 | 30.25 | -7.41 | 37.66 |  |
| Eudorcas\_thomsonii\_BOVIDAE\_CETARTIODACTYLA | Endurer | D | ? | H | 23 | 2640[6] | Semi-Arid | 0.46 | 20.63 | 28.89 | 12.61 | 16.28 |  |
| Felis\_catus\_FELIDAE\_CARNIVORA | Evaporator | D | ? | C | 5 | 3250[1] | Semi-Arid | 0.36 | 20.08 | 33.98 | 5.87 | 28.11 |  |
| Gerbilliscus\_leucogaster\_MURIDAE\_RODENTIA | Evader | D | V | G | 0.13 | 7767[1] | Semi-Arid | 0.49 | 21.5 | 31.7 | 8.81 | 22.9 | 1.02[2,3] |
| Gerbillurus\_paeba\_MURIDAE\_RODENTIA | Evader | DF | F | G | 0.026 | 6997[7] | Arid | 0.2 | 19.99 | 32.46 | 4.92 | 27.54 | 1.03[2,3] |
| Gerbillurus\_setzeri\_MURIDAE\_RODENTIA | Evader | DF | F | G | 0.038 | 5648[7] | Arid | 0.04 | 20.24 | 28.24 | 11.18 | 17.06 | 0.8[2,3] |
| Gerbillurus\_tytonis\_MURIDAE\_RODENTIA | Evader | D | V | G | 0.029 | 6324[1] | Arid | 0.03 | 18.64 | 29.16 | 7.81 | 21.36 | 1.06[2,3] |
| Gerbillurus\_vallinus\_MURIDAE\_RODENTIA | Evader | DF | F | G | 0.035 | 7144[7] | Arid | 0.09 | 19.05 | 33.26 | 3.68 | 29.57 | 0.9[2,3] |
| Gerbillus\_gerbillus\_MURIDAE\_RODENTIA | Evader | x | ? | G | 0.077 | 5500[4] | Arid | 0.03 | 24.68 | 39.93 | 8.01 | 31.92 | 1.06[2,3] |
| Glaucomys\_volans\_SCIURIDAE\_RODENTIA | Evader | D | F | O | 0.75 | 3200[1] | Humid | 0.97 | 12.75 | 30.07 | -5.14 | 35.21 | 1.2[2,3] |
| Hemiechinus\_auritus\_ERINACEIDAE\_EULIPOTYPHLA | Evader | P | F | O | 0.301 | 4010[4], | Semi-Arid | 0.24 | 8.48 | 30.96 | -13.47 | 44.43 | 0.38[2,3] |
| Isoodon\_macrourus\_PERAMELIDAE\_PERAMELEMORPHIA | Evaporator | D | F | I | 1.551 | 3126[1] | Dry sub-humid | 0.63 | 23 | 33.24 | 10.86 | 22.38 | 0.37[2,3] |
| Jaculus\_jaculus\_DIPODIDAE\_RODENTIA | Evader | x | ? | G | 0.06 | 6500[1] | Arid | 0.05 | 23.76 | 39.42 | 7.41 | 32 | 1.23[2,3] |
| Kobus\_ellipsiprymnus\_BOVIDAE\_CETARTIODACTYLA | Endurer | D | ? | H | 100 | 1098[1] | Dry sub-humid | 0.6 | 24.04 | 33.54 | 14.14 | 19.41 | 0.27[2,3] |
| Kobus\_kob\_BOVIDAE\_CETARTIODACTYLA | Endurer | D | ? | H | 150 | 1594[4] | Dry sub-humid | 0.62 | 26.25 | 36.04 | 16.91 | 19.14 |  |
| Lasionycteris\_noctivagans\_VESPERTILIONIDAE\_CHIROPTERA | Evader | D | F | I | 0.01 | 4325[1] | Humid | 0.83 | 7.47 | 27.3 | -11.76 | 39.06 |  |
| Lepus\_californicus\_LEPORIDAE\_LAGOMORPHA | Evaporator | DF | V | H | 2.4 | 3700[8] | Semi-Arid | 0.39 | 13.8 | 32.54 | -3.48 | 36.02 | 0.57[2,3] |
| Lepus\_capensis\_LEPORIDAE\_LAGOMORPHA | Evaporator | S | V | H | 1.4 | 4470[9] | Arid | 0.17 | 22.42 | 36.38 | 7.76 | 28.63 |  |
| Liomys\_irroratus\_HETEROMYIDAE\_RODENTIA | Evader | D | V | G | 0.068 | 4280[1] | Dry sub-humid | 0.5 | 19.26 | 31.13 | 6.42 | 24.71 | 1.12[2,3] |
| Liomys\_pictus\_HETEROMYIDAE\_RODENTIA | Evader | D | ? | G | 0.04 | 3950[1] | Dry sub-humid | 0.65 | 22.8 | 33.88 | 11.16 | 22.72 |  |
| Macaca\_mulatta\_CERCOPITHECIDAE\_PRIMATES | Evaporator | D | F | O | 3 | 1607[1] | Humid | 0.88 | 19.08 | 32.21 | 3.96 | 28.25 |  |
| Macropus\_eugenii\_MACROPODIDAE\_DIPROTODONTIA | Evaporator | SP | F | H | 3 | 3301[1] | Semi-Arid | 0.48 | 15.82 | 27.52 | 6.69 | 20.84 | 0.29[2,3] |
| Macropus\_robustus\_MACROPODIDAE\_DIPROTODONTIA | Endurer | D | F | H | 19.8 | 4054[1] | Semi-Arid | 0.26 | 22.57 | 36.38 | 7.69 | 28.69 | 0.19[2,3] |
| Macropus\_rufus\_MACROPODIDAE\_DIPROTODONTIA | Endurer | D | F | H | 18.7 | 3270[1] | Arid | 0.18 | 22.51 | 36.99 | 7.13 | 29.86 | 0.18[2,3] |
| Macrotis\_lagotis\_THYLACOMYIDAE\_PERAMELEMORPHIA | Evaporator | D | F | CI | 1.011 | 3976[1] | Arid | 0.16 | 25.53 | 39.29 | 9.32 | 29.97 | 0.35[2,3] |
| Madoqua\_kirkii\_BOVIDAE\_CETARTIODACTYLA | Evaporator | D | ? | H | 4.5 | 4762[1] | Semi-Arid | 0.36 | 22.38 | 30.92 | 13.18 | 17.74 | 0.5[2,3] |
| Meriones\_unguiculatus\_MURIDAE\_RODENTIA | Evader | D | F | G | 0.07 | 4958[4] | Semi-Arid | 0.43 | 1.43 | 25.02 | -23.97 | 48.99 | 1.15[2,3] |
| Mesocricetus\_auratus\_CRICETIDAE\_RODENTIA | Evader | D | F | O | 0.105 | 5340[1] | Semi-Arid | 0.24 | 17.26 | 37.73 | 0.42 | 37.31 | 1.5[2,3] |
| Microtus\_ochrogaster\_CRICETIDAE\_RODENTIA | Evader | D | F | H | 0.04 | 2362[1] | Dry sub-humid | 0.65 | 7.97 | 28.98 | -13.34 | 42.32 | 1.749[2,3] |
| Microtus\_pennsylvanicus\_CRICETIDAE\_RODENTIA | Evader | D | F | H | 0.05 | 1663[1] | Humid | 0.97 | -0.2 | 21.97 | -22.05 | 44.02 | 2.65[2,3] |
| Myodes\_gapperi\_CRICETIDAE\_RODENTIA | Evader | D | F | H | 0.019 | 1893[1] | Humid | 1.02 | 1.19 | 22.76 | -20.69 | 43.45 | 1.75[2,3] |
| Myotis\_auriculus\_VESPERTILIONIDAE\_CHIROPTERA | Evader | D | F | I | 0.007 | 3700[1] | Semi-Arid | 0.38 | 17.26 | 32.71 | 1.43 | 31.27 |  |
| Myotis\_lucifugus\_VESPERTILIONIDAE\_CHIROPTERA | Evader | D | F | I | 0.007 | 3700[1] | Humid | 0.97 | 3.15 | 24.09 | -17.51 | 41.6 | 1.43[2,3] |
| Myotis\_thysanodes\_VESPERTILIONIDAE\_CHIROPTERA | Evader | D | F | I | 0.006 | 3800[1] | Semi-Arid | 0.42 | 12.85 | 31.22 | -3.21 | 34.43 |  |
| Myotis\_volans\_VESPERTILIONIDAE\_CHIROPTERA | Evader | D | ? | I | 0.006 | 3525[1] | Dry sub-humid | 0.64 | 8.45 | 27.56 | -8.04 | 35.59 |  |
| Nanger\_granti\_BOVIDAE\_CETARTIODACTYLA | Endurer | D | ? | H | 50 | 2789[1] | Semi-Arid | 0.32 | 24.19 | 32.23 | 16.78 | 15.45 |  |
| Neovison\_vison\_MUSTELIDAE\_CARNIVORA | Evaporator | S | F | C | 1.6 | 2087[4] | Humid | 0.94 | 1.48 | 23.05 | -19.48 | 42.53 | 0.74[2,3] |
| Notomys\_alexis\_MURIDAE\_RODENTIA | Evader | D | FV | G | 0.029 | 9374[4] | Arid | 0.14 | 22.77 | 37.88 | 6.88 | 30.99 | 1.4[2,3] |
| Notomys\_cervinus \_MURIDAE\_RODENTIA | Evader | S | FV | G | 0.035 | 5510[1] | Arid | 0.11 | 23.8 | 38.76 | 7.16 | 31.6 | 1.33[2,3] |
| Nycticebus\_coucang\_LORISIDAE\_PRIMATES | Evaporator | D | ? | O | 1.38 | 3000[1] | Humid | 1.58 | 25.33 | 30.45 | 20.64 | 9.82 | 0.24[2,3] |
| Ochrotomys\_nuttalli\_CRICETIDAE\_RODENTIA | Evader | D | F | G | 0.031 | 3166[1] | Humid | 0.97 | 15.92 | 32.05 | -0.87 | 32.92 | 1.39[2,3] |
| Octomys\_mimax\_OCTODONTIDAE\_RODENTIA | Evader | x | V | H | 0.09832 | 2071[5] | Arid | 0.15 | 15.97 | 30.18 | 0.72 | 29.47 | 0.97[2,3] |
| Ondatra\_zibethicus\_CRICETIDAE\_RODENTIA | Evaporator | D | F | H | 1.124 | 1063[1] | Humid | 0.86 | 0.94 | 23.67 | -21.06 | 44.73 | 0.64[2,3] |
| Onychomys\_torridus\_CRICETIDAE\_RODENTIA | Evader | DPS | F | CI | 0.026 | 4250[1] | Semi-Arid | 0.21 | 16.87 | 35.89 | 0.21 | 35.67 | 1.55[2,3] |
| Oryctolagus\_cuniculus\_LEPORIDAE\_LAGOMORPHA | Evaporator | D | F | H | 1.8 | 1390[4] | Humid | 1.01 | 10.06 | 24.1 | -1.01 | 25.11 | 0.57[2,3] |
| Oryx\_beisa\_BOVIDAE\_CETARTIODACTYLA | Endurer | D | ? | H | 79 | 3100[6] | Semi-Arid | 0.39 | 23.2 | 31.06 | 15.83 | 15.24 |  |
| Oryx\_gazella\_BOVIDAE\_CETARTIODACTYLA | Endurer | x | ? | H | 220 | 2900[6] | Arid | 0.18 | 20.21 | 32.88 | 4.68 | 28.19 |  |
| Oryx\_leucoryx\_BOVIDAE\_CETARTIODACTYLA | Endurer | D | F | H | 92.5 | 2504[10] | Arid | 0.04 | 27.47 | 41.97 | 11.21 | 30.76 | 0.19[2,3] |
| Otolemur\_crassicaudatus\_GALAGIDAE\_PRIMATES | Evaporator | D | F | O | 1.5 | 2000[1] | Dry sub-humid | 0.62 | 22.08 | 30.91 | 11.65 | 19.26 | 0.43[2,3] |
| Otospermophilus\_beecheyi\_SCIURIDAE\_RODENTIA | Evader | D | V | O | 0.318 | 3300[1] | Humid | 0.74 | 12.26 | 29.93 | -0.55 | 30.48 | 0.53[2,3] |
| Paraechinus\_aethiopicus\_ERINACEIDAE\_EULIPOTYPHLA | Evader | P | F | O | 0.352 | 3634[4] | Arid | 0.07 | 23.96 | 39.9 | 7.88 | 32.02 | 0.25[2,3] |
| Perodicticus\_potto\_LORISIDAE\_PRIMATES | Evaporator | D | F | O | 1.125 | 1935[1] | Humid | 1.05 | 25.9 | 33.18 | 19.7 | 13.48 | 0.37[2,3] |
| Peromyscus\_polionotus\_CRICETIDAE\_RODENTIA | Evader | D | F | O | 0.014 | 3240[1] | Humid | 0.95 | 17.8 | 32.66 | 2.03 | 30.63 | 1.17[2,3] |
| Procavia\_capensis\_PROCAVIIDAE\_HYRACOIDEA | Evaporator | D | F | H | 2.4 | 3200[4] | Semi-Arid | 0.34 | 24.78 | 35.94 | 12.84 | 23.11 | 0.4[2,3] |
| Psammomys\_obesus\_MURIDAE\_RODENTIA | Evader | D | F | H | 0.15 | 6340[1] | Arid | 0.07 | 22.09 | 38.32 | 6.02 | 32.3 |  |
| Pseudomys\_hermannsburgensis\_MURIDAE\_RODENTIA | Evader | D | F | G | 0.0126 | 8970[1] | Arid | 0.14 | 22.77 | 37.74 | 6.92 | 30.82 | 1.91[2,3] |
| Rangifer\_tarandus\_CERVIDAE\_CETARTIODACTYLA | Endurer | D | F | H | 64 | 837[1] | Humid | 1.01 | -8.42 | 17.57 | -32.09 | 49.65 | 0.31[2,3] |
| Rattus\_rattus\_MURIDAE\_RODENTIA | Evader | D | F | O | 0.172 | 2900[1] | Humid | 0.84 | 17.81 | 31.1 | 4.48 | 26.62 |  |
| Rattus\_villosissimus\_MURIDAE\_RODENTIA | Evader | D | ? | O | 0.141 | 3457[1] | Semi-Arid | 0.21 | 24.15 | 37.82 | 8.45 | 29.37 | 0.58[2,3] |
| Rhinolophus\_ferrumequinum\_RHINOLOPHIDAE\_CHIROPTERA | Evader | D | F | I | 0.015 | 2980[1] | Humid | 0.75 | 12.42 | 28.99 | -3.19 | 32.18 |  |
| Rhinopoma\_hardwickii\_RHINOPOMATIDAE\_CHIROPTERA | Evader | D | F | I | 0.013 | 3722[1] | Arid | 0.17 | 24.76 | 39.16 | 9.25 | 29.91 |  |
| Saguinus\_oedipus\_CALLITRICHIDAE\_PRIMATES | Evader | D | F | O | 0.6 | 2300[1] | Humid | 1.13 | 27.03 | 32.67 | 21.73 | 10.94 |  |
| Salinomys\_delicatus\_CRICETIDAE\_RODENTIA | Evader | x | V | O | 0.0125 | 7440[5] | Semi-Arid | 0.25 | 19.36 | 33.75 | 3.85 | 29.89 |  |
| Sapajus\_apella\_CEBIDAE\_PRIMATES | Evaporator | D | F | O | 2.2 | 1160[1] | Humid | 1.22 | 25.85 | 32.66 | 19.26 | 13.4 |  |
| Sciurus\_anomalus\_SCIURIDAE\_RODENTIA | Evader | x | ? | O | 0.265 | 1031[1] | Dry sub-humid | 0.61 | 12.5 | 30.23 | -2.51 | 32.74 |  |
| Sciurus\_carolinensis\_SCIURIDAE\_RODENTIA | Evader | D | F | O | 0.55 | 2450[1] | Humid | 0.91 | 10.78 | 29.42 | -8.66 | 38.08 | 0.84[2,3] |
| Setonix\_brachyurus\_MACROPODIDAE\_DIPROTODONTIA | Evaporator | DS | F | H | 5 | 2188[1] | Humid | 0.84 | 15.44 | 26.63 | 7.02 | 19.61 | 0.3[2,3] |
| Sigmodon\_hispidus\_CRICETIDAE\_RODENTIA | Evader | D | F | O | 0.136 | 2734[1] | Humid | 0.7 | 16.57 | 33.45 | -1.06 | 34.51 | 1.654[2,3] |
| Sminthopsis\_crassicaudata\_DASYURIDAE\_DASYUROMORPHIA | Evader | D | F | CI | 0.013 | 3519[4] | Arid | 0.2 | 20.21 | 35.18 | 5.51 | 29.67 | 1.29[2,3] |
| Sylvilagus\_aquaticus\_LEPORIDAE\_LAGOMORPHA | Evaporator | S | F | H | 1.8 | 2344[1] | Humid | 0.96 | 17.09 | 33.05 | 0.31 | 32.74 |  |
| Sylvilagus\_audubonii\_LEPORIDAE\_LAGOMORPHA | Evaporator | S | F | H | 0.9 | 3349[1] | Semi-Arid | 0.31 | 13.85 | 32.58 | -3.78 | 36.36 | 0.65[2,3] |
| Sylvilagus\_floridanus\_LEPORIDAE\_LAGOMORPHA | Evaporator | S | F | H | 0.9 | 2914[1] | Humid | 0.78 | 14.29 | 30.99 | -2.81 | 33.8 |  |
| Syncerus\_caffer\_BOVIDAE\_CETARTIODACTYLA | Endurer | D | ? | H | 353 | 1124[4] | Humid | 0.73 | 23.47 | 32.19 | 14.54 | 17.65 |  |
| Tachyglossus\_aculeatus\_TACHYGLOSSIDAE\_MONOTREMATA | Evaporator | D | F | I | 4.25 | 2300[1] | Semi-Arid | 0.31 | 21.78 | 35.39 | 7.54 | 27.85 | 0.13[2,3] |
| Tamiasciurus\_hudsonicus\_SCIURIDAE\_RODENTIA | Evader | D | F | O | 0.36 | 2513[1] | Humid | 1 | 0.03 | 21.99 | -21.71 | 43.7 | 1.12[2,3] |
| Tragelaphus\_oryx\_BOVIDAE\_CETARTIODACTYLA | Endurer | D | F | H | 150 | 1881[1] | Semi-Arid | 0.48 | 21.24 | 31.5 | 8.96 | 22.55 | 0.24[2,3] |
| Tragelaphus\_scriptus\_BOVIDAE\_CETARTIODACTYLA | Endurer | D | ? | H | 50 | 1369[1] | Dry sub-humid | 0.65 | 23.75 | 32.92 | 13.99 | 18.93 |  |
| Trichosurus\_vulpecula\_PHALANGERIDAE\_DIPROTODONTIA | Evaporator | D | F | H | 2.25 | 1504[1] | Dry sub-humid | 0.52 | 20.45 | 32.64 | 7.64 | 25.01 | 0.32[2,3] |
| Tupaia\_glis\_TUPAIIDAE\_SCANDENTIA | Evader | D | F | O | 0.15 | 2480[4] | Humid | 1.51 | 26.07 | 31.58 | 21.17 | 10.41 | 0.76[2,3] |
| Tympanoctomys\_barrerae\_OCTODONTIDAE\_RODENTIA | Evader | x | V | H | 0.0868 | 7080[5) | Arid | 0.19 | 14.61 | 31.58 | -0.7 | 32.29 | 1.08[2,3] |
| Urocitellus\_columbianus\_SCIURIDAE\_RODENTIA | Evader | D | FV | O | 0.5 | 2792[1] | Humid | 0.86 | 3.7 | 23.25 | -11.57 | 34.82 |  |
| Vulpes\_zerda\_CANIDAE\_CARNIVORA | Evaporator | D | F | O | 1.2 | 4022[1] | Arid | 0.02 | 25 | 40.3 | 8.14 | 32.16 | 0.48[2,3] |
| Xerospermophilus\_tereticaudus\_SCIURIDAE\_RODENTIA | Evader | D | F | H | 0.125 | 5030[1] | Arid | 0.13 | 20.25 | 38.84 | 3.22 | 35.62 | 0.56[2,3] |
