## Supplemental Figures for "Convergent evolution of increased urine concentrating ability in desert mammals"

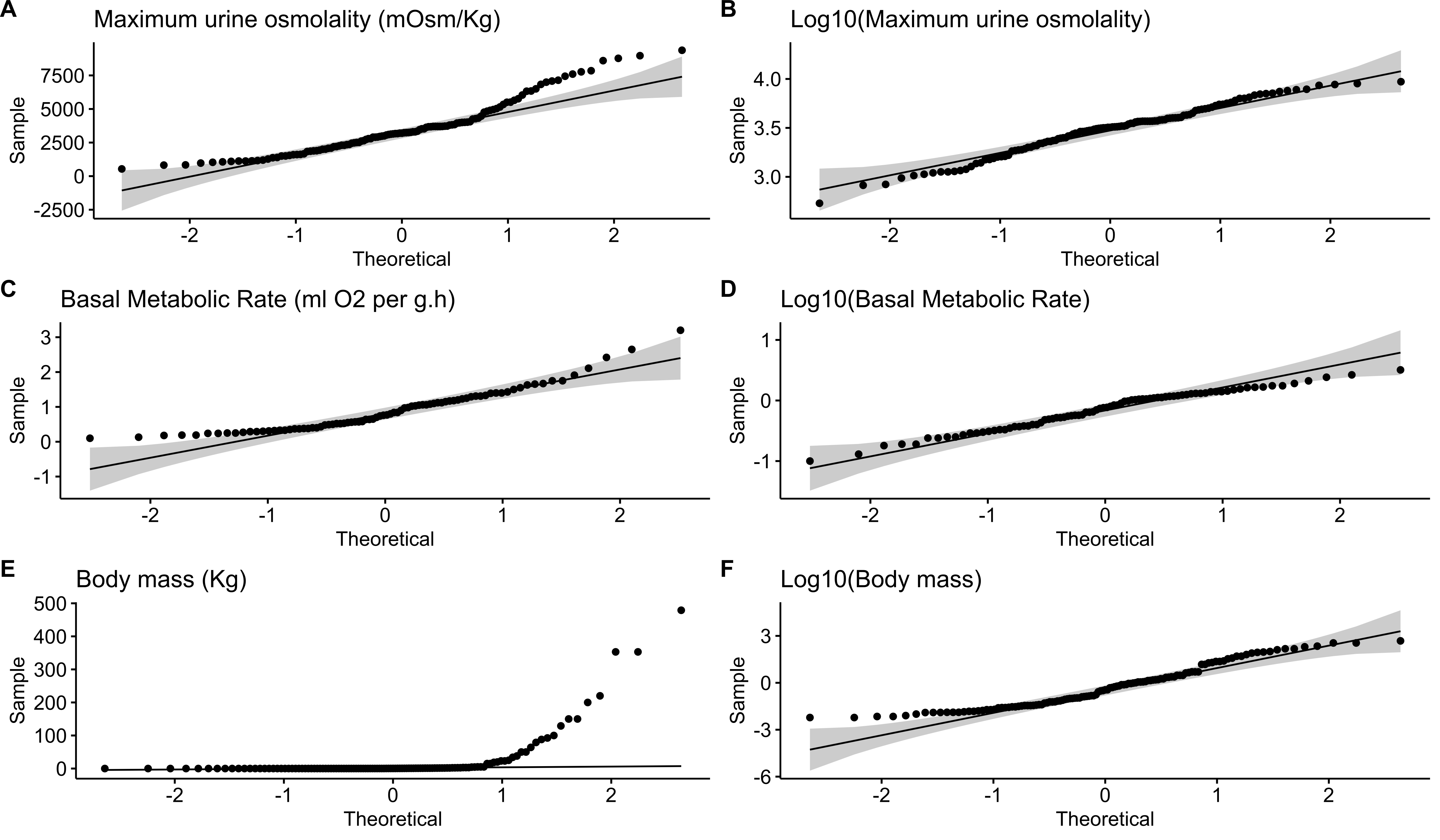

**Figure S1**– QQ plots for the normal distribution of variables (A, C, E) and log10 of variables (B, D, F) used in this study.

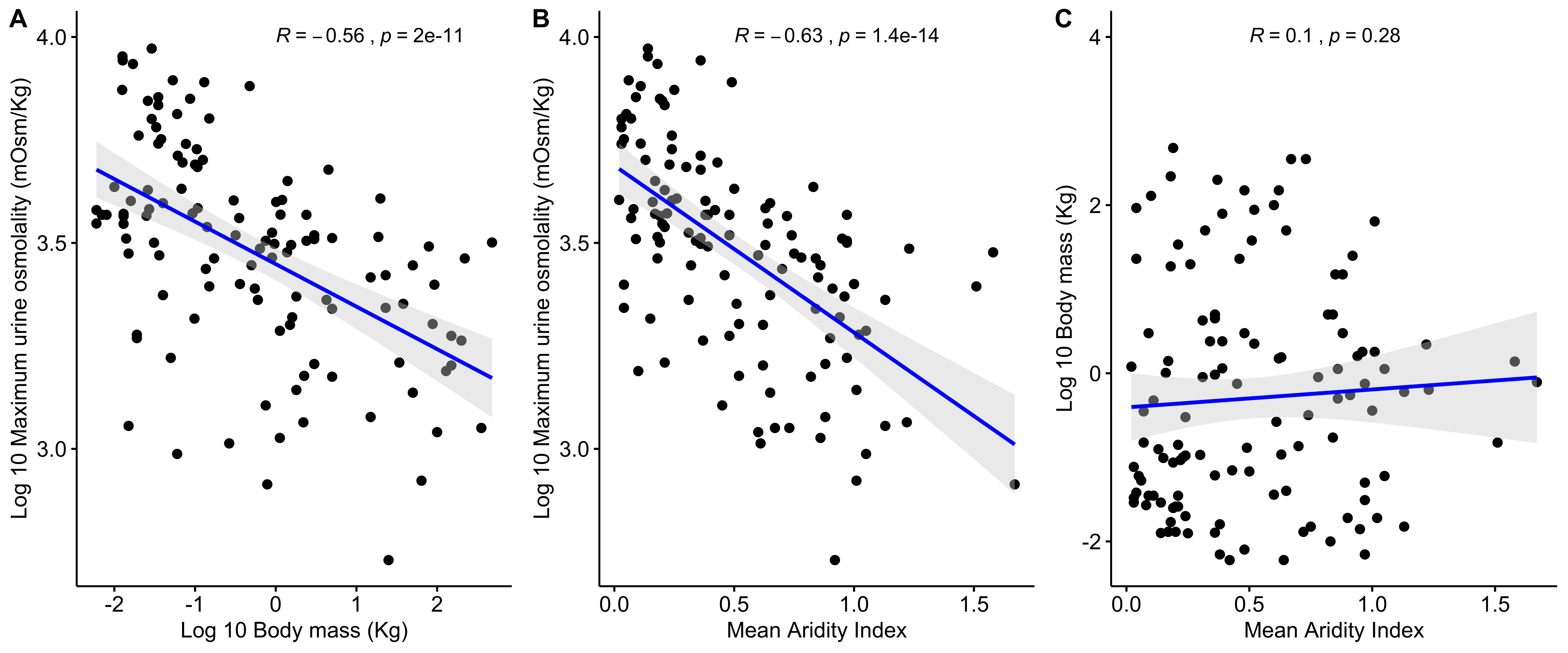

**Figure S2** – Spearman correlation for: A- Log 10 maximum urine osmolality (measured in mOsm/Kg) and Log10 body mass (Kg); B- Log10 maximum urine osmolality (measured in mOsm/Kg) and mean aridity index; and C- Log10 body mass (Kg) and mean aridity index. P-values are not presented as the plots do not take phylogenetic affinity into account.

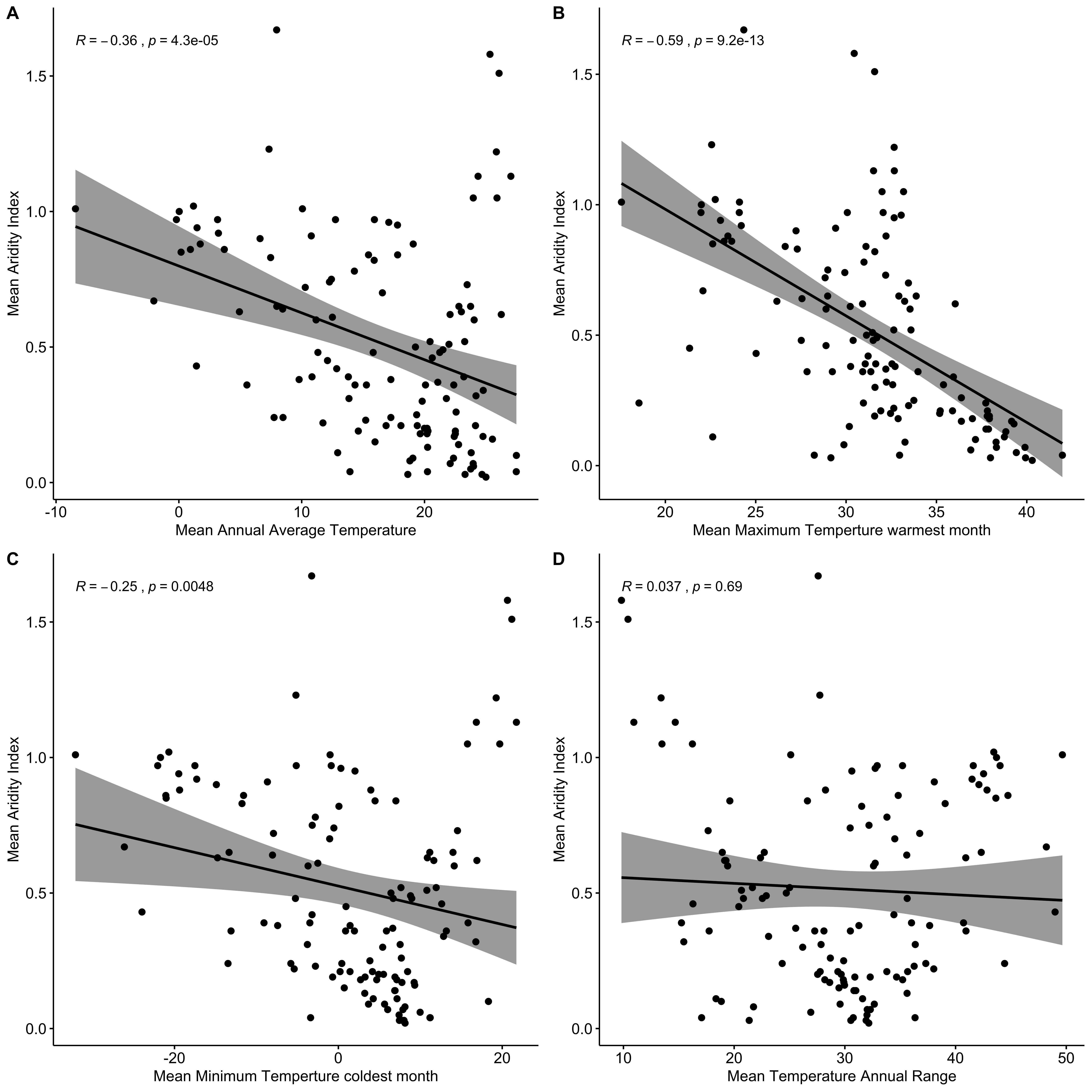

**Figure S3**– Spearman correlation for mean annual aridity index and four measurements of temperature variation. P-values are not presented as the plots do not take spatial correlation into account.

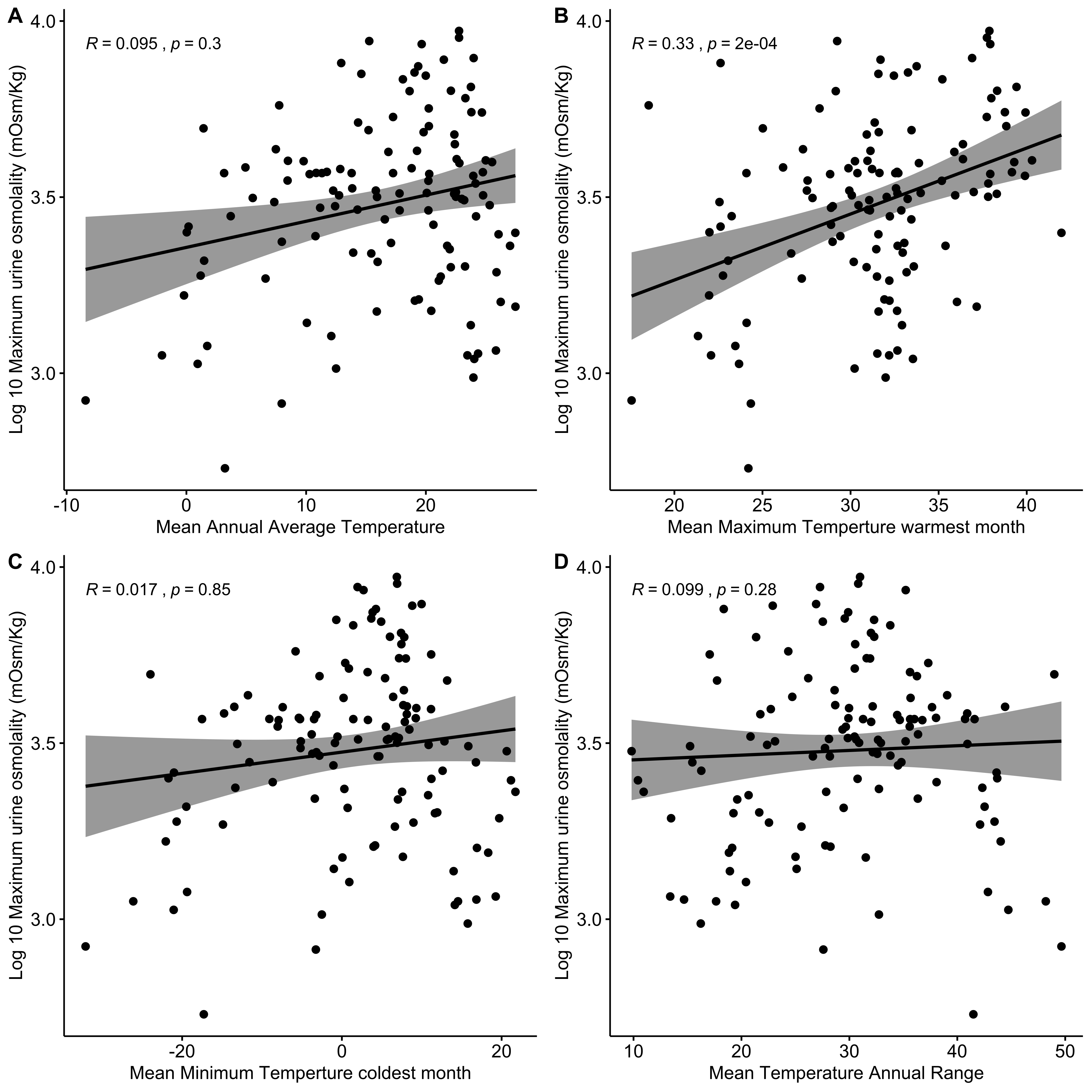

**Figure S4** – Spearman correlation for log10 maximum urine osmolality (measured in mOsm/Kg) and four measurements of temperature variation. P-values are not represented as the plots do not take phylogenetic affinity into account.

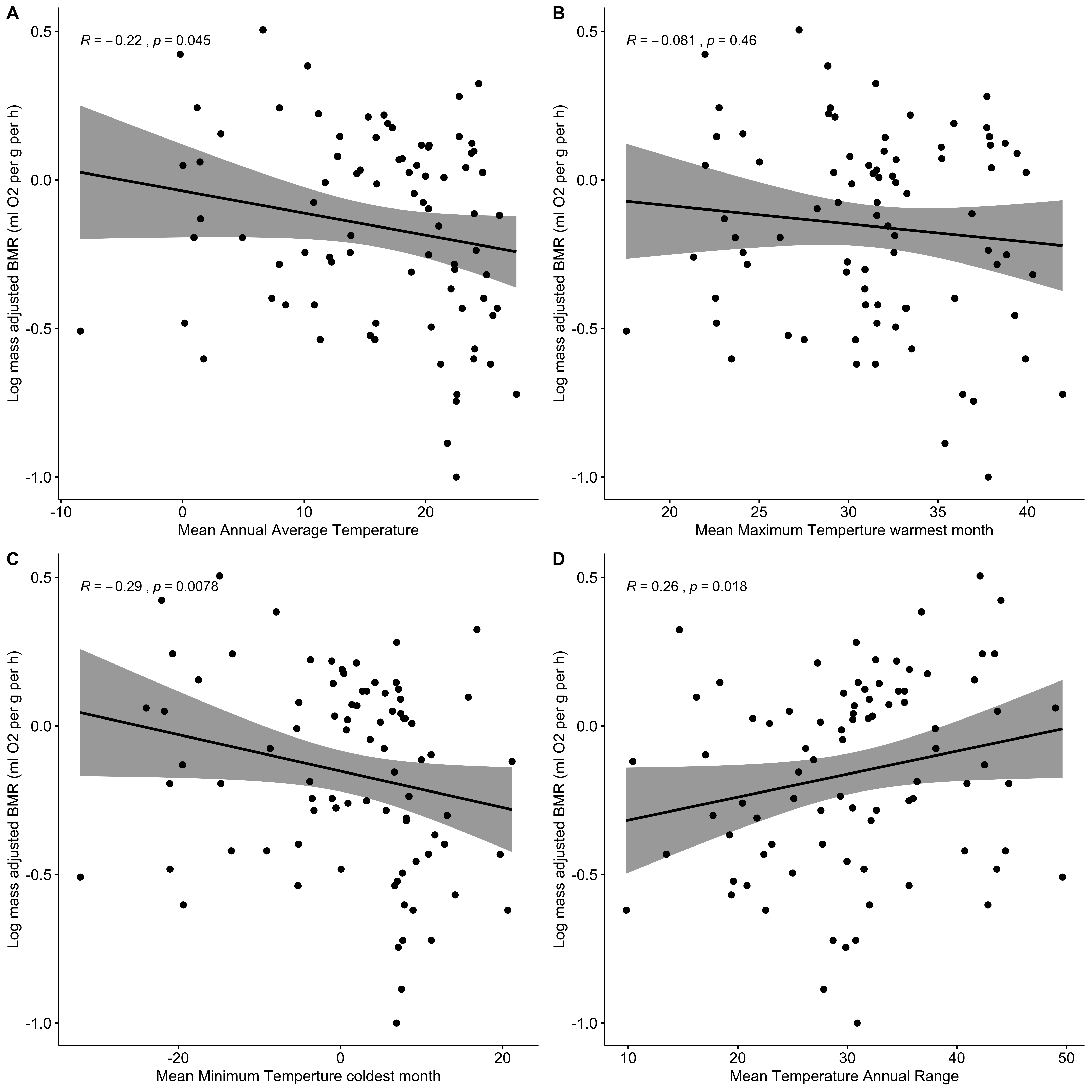

**Figure S5** – Spearman correlation for log10 mass-adjusted Basal Metabolic Rate (ml of O_2_ per g per day) and four measurements of temperature variation. P-values are not represented as plots do not take phylogenetic affinity into account.
