## Supplemental Tables S2-S4 for "Convergent evolution of increased urine concentrating ability in desert mammals": Rocha_et_al_SupplementaryMaterial_TableS2-S4.html

**Table S2 –
PGLS models for predicting log10 mammalian maximum urine osmolality (mOsm/kg)
ordered by increasing complexity.** Variables are
AI = mean annual aridity index (continuous), cAI = mean annual aridity index
(categorical: Arid, Semi-Arid, Dry sub-humid, Humid), BM = Log10 body mass
(kg), C = condition  (dehydrated-D
relative to fed-F, given salt load-S, given protein
load-P, unknown-x and combinations of these), M
= Method (unknown-? relative to addition of solutes-A, Freezing point-F, Vapor
pressure-V, and combinations of these), Di = Diet (Carnivorous-C relative to
Fructivorous-Fr, Granivorous-G, Herbivorous-H, Insectivorous-I, Omnivorous-O,
and combinations of these), BIO1 = mean Annual Mean Temperature, BIO5= mean
Maximum Temperature of  Warmest Month, BIO6
= mean Minimum Temperature of Coldest Month, BIO7 = mean Temperature Annual
Range. Highlighted are significant variables (p-value < 0.05). When **λ** is zero the covariance between species is zero and the
linear model approaches that of a non-phylogenetic regression, while when **λ**
is 1 the evolution of the residual error follows a Brownian model.

|  |  |  |  |
| --- | --- | --- | --- |
| Dataset | Model | AIC | Pagel’s **λ** |
| Different conditions  (121 species) | **AI** | -60.17 | 0.8561328 |
|  | **BIO1** | -35.19 | 0.9748651 |
|  | **BIO5** | -37.42 | 0.9680426 |
|  | **BIO6** | -30.00 | 0.9796061 |
|  | BIO7 | -22.21 | 0.9845628 |
|  | **AI** + **BM** | -74.36 | 0.2219936 |
|  | **AI** + **BM** + C | -38.55 | 0.158529 |
|  | **AI + BM** + M | -54.64 | 0.1146517 |
|  | **AI** + **BM** + **Di** | -66.33 | -0.0146275 |
|  | **AI + BM** + C + M | -19.12 | 0.08665056 |
|  | **AI + BM** + C + D | -29.92 | -0.01394202 |
|  | **AI + BM** + C + M + **Di** | -16.29 | -0.02352754 |
|  | **BIO1** + **BM** | -45.61 | 0.9632151 |
|  | **BIO1** + **BM** + C | -11.50 | 0.9658072 |
|  | **BIO1** + **BM** + M | -26.52 | 0.9606075 |
|  | **BIO1** + **BM** + **Di** | -31.33 | 0.9043004 |
|  | **BIO1** + **BM** + C + M + **Di** | 18.32 | 0.8811903 |
|  | **BIO5** + **BM** | -49.39 | 0.955564 |
|  | **BIO5** + **BM** + C | -14.68 | 0.9561875 |
|  | **BIO5** + **BM** + M | -30.89 | 0.9517756 |
|  | **BIO5** + **BM** + **Di** | -34.38 | 0.8907402 |
|  | **BIO5** + **BM** + C + M + **Di** | 12.57 | 0.0488838 |
|  | **BIO6** + **BM** | -39.56 | 0.9679175 |
|  | **BIO6** + **BM** + C | -6.72 | 0.9712682 |
|  | **BIO6** + **BM** + M | -20.75 | 0.9655426 |
|  | **BIO6** + **BM** + **Di** | -24.22 | 0.9170073 |
|  | **BIO6** + **BM** + C + M + Di | 24.33 | 0.9169933 |
| Dehydrated only  (87 species) | **AI** | -54.40 | 0.9436382 |
|  | **BIO1** | -34.04 | 0.9969119 |
|  | **BIO5** | -32.34 | 0.9876165 |
|  | **BIO6** | -32.31 | 1.001864 |
|  | **BIO7** | -28.63 | 1.004338 |
|  | **AI** + **BM** | -62.28 | 0.9294967 |
|  | **AI** + **BM** + M | -42.76 | 0.898354 |
|  | **AI** + **BM** + **Di** | -52.34 | -0.03254744 |
|  | **AI** + **BM** + M + **Di** | -34.72 | -0.02274544 |
|  | **AI** + **BM** + M + **Di** + BIO7 | -23.86 | -0.03390474 |
|  | **BIO1** + **BM** | -40.68 | 0.99556 |
|  | **BIO1** + **BM** + M | -20.44 | 0.995414 |
|  | **BIO1** + **BM** + **Di** | -27.48 | 0.9877439 |
|  | **BIO1** + **BM** + M + **Di** | -8.06 | 0.9871385 |
|  | **BIO5** + **BM** | -39.52 | 0.985862 |
|  | **BIO5**+ **BM** + M | -20.58 | 0.9851374 |
|  | **BIO5** + **BM** + **Di** | -26.56 | 0.9676973 |
|  | **BIO5** + **BM** + M + **Di** | -5.79 | -0.02165534 |
|  | **BIO6** + **BM** | -37.98 | 1.000221 |
|  | **BIO6**+ **BM** + M | -17.50 | 0.999279 |
|  | **BIO6** + **BM** + **Di** | -23.94 | 0.9957233 |
|  | **BIO6** + **BM** + M + **Di** | -4.36 | 0.9942829 |
|  | **BIO7** + **BM** | -33.44 | 1.002295 |
|  | **BIO7**+ **BM** + M | -13.35 | 1.001486 |
|  | **BIO7**+ **BM** + **Di** | -17.57 | 0.9999706 |
|  | **BIO7**+ **BM** + M+ **Di** | 1.47 | 0.9991637 |
|  | **cAI** | -42.76 | 0.9528914 |
|  | **cAI** + **BM** | -48.88 | 0.9504767 |
|  | **cAI** + **BM** + M | -29.49 | 0.9392195 |
|  | **cAI** + **BM** + **Di** | -42.68 | -0.03279375 |
|  | **cAI** + **BM** + M + **Di** | -24.99 | -0.01932766 |
|  | **cAI** + **BM** + M + D + BIO7 | -10.78 | -0.0192356 |

**Table S3
–PGLS models for predicting mammalian log10 maximum urine osmolality (mOsm/kg)** Model
notation refers to the combination of variables used as predictors while taking
phylogeny into account. Highlighted in bold are significant p-values. Variables
are cAI = categorical mean annual aridity index (Arid relative to Semi-Arid,
Dry sub-humid, Humid), BM = log10 body mass (kg), C = condition (dehydrated-D
relative to fed-F, given salt load-S, given protein
load-P, unknown-x and combinations of these), M = Method (unknown-? relative to
addition of solutes-A, Freezing point-F, Vapor pressure-V, and combinations of
these), Di = Diet (Carnivorous-C relative to Fructivorous-Fr, Granivorous-G,
Herbivorous-H, Insectivorous-I, Omnivorous-O, and combinations of these).

|  |  |  |  |  |  |  |  |
| --- | --- | --- | --- | --- | --- | --- | --- |
| *Dataset* | *Model* | *AIC* | *variable* | *Coefficient* | *s.e* | *t-value* | *p-value* |
| *Dehydrated* | cAI + BM + M +Di | -24.99 | intercept | 3.663 | 0.137 | 26.71 | **0.0000** |
|  |  |  | Dry sub-humid | -0.242 | 0.051 | -4.72 | **0.0000** |
|  |  |  | Humid | -0.342 | 0.045 | -7.58 | **0.0000** |
|  |  |  | Semi-Arid | -0.105 | 0.045 | -2.37 | **0.0207** |
|  |  |  | BM | -0.068 | 0.016 | -4.32 | **0.0000** |
|  |  |  | A _method_ | -0.154 | 0.143 | -1.08 | 0.2829 |
|  |  |  | F _method_ | -0.035 | 0.042 | -0.83 | 0.4094 |
|  |  |  | FV _method_ | 0.084 | 0.105 | 0.80 | 0.4246 |
|  |  |  | V _method_ | 0.034 | 0.068 | 0.50 | 0.6208 |
|  |  |  | CI _diet_ | -0.116 | 0.167 | -0.70 | 0.4880 |
|  |  |  | Fr _diet_ | -0.438 | 0.179 | -2.45 | **0.0166** |
|  |  |  | G _diet_ | 0.105 | 0.144 | 0.73 | 0.4653 |
|  |  |  | H _diet_ | -0.083 | 0.137 | -0.61 | 0.5454 |
|  |  |  | I _diet_ | -0.011 | 0.143 | -0.08 | 0.9384 |
|  |  |  | O _diet_ | 0.036 | 0.140 | 0.26 | 0.7971 |

**Table S4 –
PGLS models for predicting log 10 mass-adjusted basal metabolic rate (ml of O_2_ per g per day) ordered by increasing complexity.** Variables
are BM = log10 body mass (g),  AI = mean
annual aridity index (continuous), cAI = mean annual aridity index (Arid,
Semi-Arid, Dry sub-humid, Humid), Di = Diet
(Carnivorous-C relative to Fructivorous-Fr, Granivorous-G, Herbivorous-H,
Insectivorous-I, Omnivorous-O, and combinations of these), BIO1 = mean Annual
Mean Temperature, BIO5= mean Maximum Temperature of Warmest Month, BIO6 = mean Minimum
Temperature of Coldest Month, BIO7 = mean Temperature Annual Range. Highlighted
are significant variables (p-value < 0.05).

|  |  |  |  |
| --- | --- | --- | --- |
| Dataset | Model | AIC | Pagel’s **λ** |
|  | **BM** | -27.28 | 0.7601046 |
|  | **BM** + AI | -23.61 | 0.7503346 |
|  | **BM +** AI + Di | -1.68 | 0.6288732 |
|  | **BM** **+ BIO1** + Di | 1.73 | 0.5731028 |
|  | **BM** **+** BIO5 + Di | 1.77 | 0.6274593 |
|  | **BM** **+ BIO6** + Di | 1.91 | 0.5383454 |
|  | **BM** **+** BIO7 + Di | 4.05 | 0.5487208 |
|  | **BM** **+ cAI** + Di | 5.69 | 0.5982846 |
